# Robust Population Structure Inference and Correction in the Presence of Known or Cryptic Relatedness

**DOI:** 10.1101/008276

**Authors:** Matthew P. Conomos, Mike Miller, Timothy Thornton

## Abstract

Population structure inference with genetic data has been motivated by a variety of applications in population genetics and genetic association studies. Several approaches have been proposed for the identification of genetic ancestry differences in samples where study participants are assumed to be unrelated, including principal components analysis (PCA), multi-dimensional scaling (MDS), and model-based methods for proportional ancestry estimation. Many genetic studies, however, include individuals with some degree of relatedness, and existing methods for inferring genetic ancestry fail in related samples. We present a method, PC-AiR, for robust population structure inference in the presence of known or cryptic relatedness. PC-AiR utilizes genome-screen data and an efficient algorithm to identify a diverse subset of unrelated individuals that is representative of all ancestries in the sample. The PC-AiR method directly performs PCA on the identified ancestry representative subset and then predicts components of variation for all remaining individuals based on genetic similarities. In simulation studies and in applications to real data from Phase III of the HapMap Project, we demonstrate that PC-AiR provides a substantial improvement over existing approaches for population structure inference in related samples. We also demonstrate significant efficiency gains, where a single axis of variation from PC-AiR provides better prediction of ancestry in a variety of structure settings than using ten (or more) components of variation from widely used PCA and MDS approaches. Finally, we illustrate that PC-AiR can provide improved population stratification correction over existing methods in genetic association studies with population structure and relatedness.

## Introduction

Ancestry inference with genetic data is an essential component for a variety of applications in genetic association studies, population genetics, and both personalized and medical genomics. Advances in high-throughput genotyping technology have allowed for an improved understanding of continental-level and fine-scale genetic structure of human populations, as well as other organisms. Principal components analysis (PCA) (Price et al., 2006; Patterson et al., 2006) has been the prevailing approach in recent years for both population structure inference and correction of population stratification in genome-wide association studies (GWAS) with high-density single nucleotide polymorphism (SNP) genotyping data. Other widely used methods for inference on genetic ancestry include multi-dimensional scaling (MDS) (Purcell et al., 2007), a dimension reduction method similar to PCA, and model-based methods, such as STRUCTURE (Pritchard et al., 2000), FRAPPE (Tang et al., 2005), and ADMIXTURE (Alexander et al., 2009), for proportional ancestry estimation in samples from admixed populations.

Genetic studies often include related individuals; however, most existing population structure inference methods fail in the presence of relatedness. For example, the top principal components from PCA, as well as the top dimensions from MDS, can reflect family relatedness rather than population structure when applied to samples that include relatives (Price et al., 2010). Model-based ancestry estimation methods similarly fail in the presence of relatedness as they are not able to distinguish between ancestral groups and clusters of relatives (Thornton and Bermejo, 2014). For certain family-based study designs with known pedigrees, the population structure inference method proposed by Zhu et al. (2008), where SNP loadings calculated from a PCA on pedigree founders are used to obtain principal components values for genotyped offspring, can be used. However, this approach, which we refer to as “FamPCA,” fails in the presence of cryptic or misspecified relatedness and is not applicable to most GWAS where genealogical information on sample individuals is often incomplete or unavailable. The FamPCA method requires genotype data to be available for pedigree founders, which can be prohibitive for many genetic studies. In addition, inference on population structure is limited to the ancestries in the subset of genotyped founders, which may lack sufficient diversity to be representative of the ancestries in the entire sample (Chen et al., 2013).

We address the problem of population structure inference and correction in samples with related individuals. We do not put constraints on how the individuals might be related, and we allow for the possibility that genealogical information on sample individuals could be partially or completely missing. We propose a method, which we call PC-AiR (principal components analysis in related samples), for inference on population structure from SNP genotype data in general samples with related individuals. The PC-AiR method implements a fast and efficient algorithm for the identification of a diverse subset of mutually unrelated individuals who are representative of the ancestries in the entire sample. Axes of variation are inferred using this ancestry representative subset, and coordinates along the axes are predicted for all remaining sample individuals based on genetic similarities with individuals in the ancestry representative subset. The top axes of variation (principal components) from PC-AiR are constructed to be both representative of ancestry and robust to both known or cryptic relatedness in the sample. A remarkable feature of PC-AiR is the method’s ability to identify a diverse and representative subset of individuals for ancestry inference using only genome-screen data from the sample, without requiring additional samples from external reference population panels or genealogical information on the study individuals.

We assess the robustness and accuracy of PC-AiR for inference on genetic ancestry in simulation studies with both related and unrelated individuals under various types of population structure settings, including admixture. We also directly compare PC-AiR to existing population structure inference methods using both simulated data and real genotype data collected from the Mexican Americans in Los Angeles, California (MXL) and African American individuals in the southwestern USA (ASW) population samples of release 3 of phase III of the International Haplotype Map Project (HapMap) (International HapMap 3 Consortium, 2010). The population structure inference methods to which we compare PC-AiR are: (1) PCA with the EIGENSOFT (Price et al., 2006) software, (2) MDS with the PLINK (Purcell et al., 2007) software, (3) the model-based ancestry estimation methods FRAPPE (Tang et al., 2005) and ADMIXTURE (Alexander et al., 2009), and (4) FamPCA (Zhu et al., 2008) as implemented in the KING (Manichaikul et al., 2010) software. We also perform simulation studies to assess population structure correction with PC-AiR in GWAS with relatedness and ancestry admixture. We evaluate the type-I error when using PC-AiR principal components as well as widely used population stratification correction methods including: (1) the EIGENSTRAT (Price et al., 2006) method, which uses PCA with EIGENSOFT to correct for population structure, and (2) the linear mixed model methods EMMAX (Kang et al., 2010) and GEMMA (Zhou and Stephens, 2012), which use variance components and an empirical genetic relatedness matrix to simultaneously account for both population structure and relatedness among sample individuals.

## Materials & Methods

### Overview of the PC-AiR Method

Let the set *𝒩* be a sample of outbred individuals who have been genotyped in a genome-screen. An essential component of the PC-AiR method for population structure inference in the presence of relatedness is to use genome-screen data to partition *𝒩* into two non-overlapping subsets, *𝒰* and *ℛ*, i.e. *𝒩 = 𝒰* ∪ *ℛ* with *𝒰* ∩ *ℛ* = Ø, where *𝒰* is a subset of mutually unrelated individuals who are representative of the ancestries of all individuals in *𝒩*, and *ℛ* is a “related subset” of individuals who have at least one relative in *𝒰*. We allow for individuals in *ℛ* to be related to each other in addition to having relatives in *𝒰*. PC-AiR uses measures of pairwise relatedness and ancestry divergence calculated from autosomal SNP genotype data for the identification of *𝒰*, without requiring external reference panels or genealogical information. Population structure inference on the entire set of sample individuals, *𝒩*, is then obtained by first directly performing PCA on the selected ancestry representative subset, *𝒰*, and then predicting values along the components of variation for all individuals in the related subset, *ℛ*, based on genetic similarities with the individuals in *𝒰*. In the following subsections, we describe the PC-AiR method in detail.

### Population Genetic Modeling Assumptions

The population genetic modeling assumptions we make are weak and are satisfied by commonly used models of population structure, such as the Balding-Nichols model (Balding and Nichols, 1995). The individuals in set *𝒩* are assumed to have been sampled from a population with ancestry derived from *K* ancestral subpopulations. Let *𝒮* be the set of autosomal SNPs in the genome-screen, and for SNP *s* ∈ *S*, denote 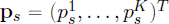 to be the vector of subpopulation-specific allele frequencies, where 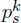 is the allele frequency at SNP *s* in subpopulation *k* ∈ {1, …, *K*}. We assume that the 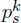 are random variables that are independent across *s* but with possible dependence across the *k*’s, with mean E[**p***_s_*] = *p_s_* **1** and covariance Cov[**p***_s_*] = *p_s_*(1−*p_s_*)**Σ***_K_* for every *s*, where **1** is a length *K* column vector of 1’s, and **Σ***_K_* is a *K × K* matrix. In genetic models incorporating population structure, the allele frequency parameter *p_s_* is typically interpreted as an “ancestral” allele frequency, or some average of allele frequencies across subpopulations. Although we allow **Σ***_K_* to be completely general, including allowing for non-zero covariances across subpopulations, a special case is the Balding-Nichols model, where **Σ***_K_* is a diagonal matrix with (*k, k*)-th element equal to *F_k_* ⩾ 0, and *F_k_* is Wright’s standardized measure of variation (Wright, 1949) for subpopulation *k*. We allow for sample individuals to have admixed ancestry from the *K* subpopulations, and we denote 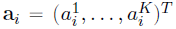 to be the ancestry vector for individual *i* ∈ *𝒩*, where 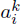 is the proportion of ancestry across the autosomal chromosomes from subpopulation *k* for individual *i*, with 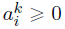 ⩾ 0 for all *k*, and 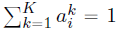. In most contexts, the parameters *K*, **Σ***_K_*, **p***_s_* and *p_s_* for all *s* ∈ *S*, and **a***_i_* for all *i* ∈ *𝒩* will be unknown. The goal of PC-AiR is to obtain inference on ancestry, i.e. the **a***_i_*’s, for all sample individuals *i* ∈ *𝒩* in the presence of known or cryptic relatedness.

### Relatedness Inference in Structured Populations

PC-AiR uses kinship coefficients to measure relatedness between all pairs of individuals in *𝒩*, where the kinship coefficient for individuals *i* and *j*, which we denote as *φ_ij_*, is defined to be the probability that a random allele selected from *i* and a random allele selected from *j* at a locus are identical-by-descent (IBD). When the genealogy of the sample individuals is known, PC-AiR can use theoretical or pedigree-based kinship coefficients, and a number of software packages (Abney, 2009; Zheng and Bourgain, 2009) are available for calculating these according to a specified genealogy. However, genealogical information on sample individual is often unknown, incomplete, or misspecified, and PC-AiR can also use empirical kinship coefficients estimated from genome-screen data for samples with cryptic relatedness that must be genetically inferred. It is important to note that relatedness estimators that assume population homogeneity, such as those implemented in the widely used PLINK software (Purcell et al., 2007) or obtained via a standard genetic relationship matrix (GRM)(Yang et al., 2010), are biased in samples from structured populations. Therefore, we do not recommended using these estimators with PC-AiR as it has been demonstrated that they give inflated kinship estimates in the presence of population structure (Thornton et al., 2012; Manichaikul et al., 2010), where (1) unrelated pairs of individuals with similar ancestry can have kinship-coefficient estimates corresponding to values that are expected for close relatives, and (2) related individuals can have a systematic inflation in their estimated degree of relatedness.

To use the PC-AiR method when pedigree relationships are unknown or incomplete, we recommend using empirical kinship coefficient estimates from methods that have been developed for samples from structured populations. One such estimator is KING (kinship-based inference for GWASs)-robust (Manichaikul et al., 2010). Rather than using estimated allele frequencies, which leads to biased relatedness estimates in the presence of population structure, KING-robust relies on shared genotype counts across the SNPs in the genome-screen to measure the genetic distance between individuals. KING-robust was developed for relatedness inference in samples from populations with discrete substructure without admixture, and it is a consistent estimator of the kinship coefficient for a pair of outbred individuals from the same subpopulation. The estimator, however, will generally be negatively biased for pairs of individuals that have different ancestries. Despite this bias, the KING-robust estimator is typically able to separate close relatives with similar ancestry from unrelated individuals, which is often sufficient for the PC-AiR method. Additionally, the PC-AiR method exploits the negative bias of the KING-robust estimator to gain insight on ancestry differences among individuals, as discussed in more detail in the following subsection.

Estimated kinship coefficients from the recently proposed REAP (Thornton et al., 2012) and RelateAdmix (Moltke and Albrechtsen, 2014) methods can also be used by PC-AiR. Both of these methods offer improved relatedness inference over KING-robust in samples with admixed ancestry by using external reference panels. REAP and RelateAdmix, however, may not be suitable for some studies as they require (1) some prior knowledge about the ancestries that are likely present in the sample, and (2) appropriate reference panels with suitable surrogates for the ancestral subpopulations. KING-robust does not require external reference panels and can be used with PC-AiR for admixed samples with cryptic relatedness when the REAP and RelateAdmix methods may not be practical.

### Measuring Ancestry Divergence with Genome-Screen Data

Pairwise measures of relatedness, such as kinship coefficients, among individuals in a sample can be used for selecting a subset of mutually unrelated individuals (Staples et al., 2013). In structured samples, however, identifying a subset of unrelated individuals based solely on relatedness measures can result in a subset that lacks sufficient diversity for population structure inference on the entire sample, as it may not be representative of the ancestries of all individuals. For the identification of an ancestry representative subset of mutually unrelated individuals, PC-AiR incorporates measures of ancestry divergence in addition to the kinship coefficients used as measures of relatedness.

Consider a pair of individuals *i, j* ∈ *𝒩* who have non-missing genotype data at the set *S_ij_* ⊂ *𝒮* of autosomal SNPs in a genome-screen, and let |*S_ij_*| denote the total number of SNPs in this set. Additionally, let the random variables *g_is_* and *g_js_*be the number of copies of the reference allele that individuals *i* and *j* each have, respectively, at SNP *s* ∈ *S_ij_*; thus, *g_is_*and *g_js_* take values of 0, 1, or 2. To measure ancestry divergence between a pair of unrelated individuals *i* and *j*, we use the estimator

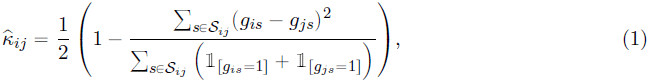

where 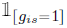 is an indicator for individual *i* being heterozygous at SNP *s*, i.e. 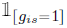 is 1 if *g_is_* = 1 and is 0 otherwise, and 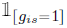 is similarly defined for individual *j*. Equation (1) is equivalent to the KING-robust estimator (Manichaikul et al., 2010) that has been proposed for estimating kinship coefficients of related individuals in samples from discrete subpopulations. We consider the KING-robust estimator under the general population genetic modeling assumptions previously discussed for *i* and *j* with admixed ancestry from *K* ancestral subpopulations. Recall that **a***_i_* and **a***_j_* are the ancestry vectors for *i* and *j*, respectively. Under an assumption that genotypes at different SNPs are independent and with |*S_ij_*| →∞, it can be shown (see the Appendix) that

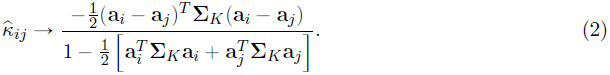

For unrelated *i* and *j* with the same ancestry, 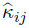 → 0, as can be seen from Equation (2) by setting **a***_i_* = **a***_j_*. However, when *i* and *j* have different ancestral backgrounds, 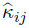 is a negatively biased estimator of kinship, and this bias provides a useful measure of the ancestry divergence between the pair of individuals. Consider, for example, the Balding-Nichols model where **Σ***_K_* is a diagonal matrix with (*k, k*)-th element equal to *F_k_*. Under this model, it can be seen from Equation (2) that

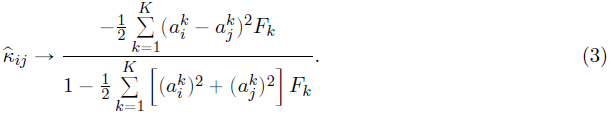

The value in Equation (3) is negative when *i* and *j* have different ancestry proportions, and the magnitude of this negative value will depend on how divergent the ancestries for the pair are. The 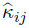 estimator will generally have more extreme negative values when (1) the *F_k_* values are large, (2) *i* and *j* have large ancestry proportion differences, or (3) either *i* or *j* has an ancestry proportion that is close to 1 from one of the *K* subpopulations. For the special case when *i* and *j* are non-admixed and have ancestry from different subpopulations *k* and *k′*, the estimator reaches an extreme negative value and

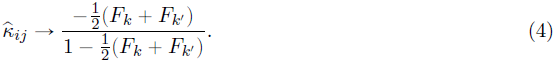

PC-AiR uses the 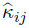 estimator given by Equation (1) for inference on ancestry divergence for all pairs of individuals *i, j* ∈ *𝒩* who are not inferred to be related based on the kinship coefficient measures discussed in the previous subsection.

### Identification of an Ancestry Representative Subset

We now provide details on how PC-AiR uses both the relatedness and ancestry divergence measures discussed in the previous two subsections for the identification of *U,* a mutually unrelated subset of individuals that is representative of the ancestries of all individuals in the sample *𝒩*. Let 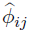 be the kinship coefficient measure that is chosen for relatedness inference on a pair of individuals *i, j* ∈ *𝒩*. When the genealogy of the sample individuals is known, 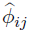 could be a pedigree-based kinship coefficient, and when the genealogy is partially or completely unknown, 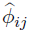 would be an empirical kinship coefficient estimate from a relatedness estimation method that allows for population structure, e.g., the KING-robust estimator of Equation (1), REAP, or RelateAdmix. In order to identify all pairs of relatives in *𝒩*, a relatedness threshold, *τ_φ_*, is chosen such that *i* and *j* are designated to be related by the PC-AiR method if 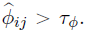. When pedigree-based kinship coefficients are used with PC-AiR, all unrelated pairs will have 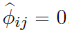, and *τ_φ_* should be set to 0. When empirical kinship coefficient estimates are used, there will be some noise in the estimation, and *τ_φ_* can be set to an approximate upper bound that is expected for the chosen kinship coefficient estimator for an unrelated pair. For example, when using KING-robust for relatedness inference, i.e. using 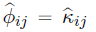, we have found that 0.025 is an approximate upper bound with dense SNP genotyping data for unrelated pairs with the same ancestry, and setting *τ_φ_* = 0.025 works well in practice for identifying relatives with similar ancestry up to third-degree (and some fourth-degree) in a variety of population structure settings with ancestry admixture. For all sample individuals *i* ∈ *𝒩*, we calculate 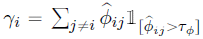 as a measure of the total kinship individual *i* has with its inferred relatives in the sample, where 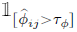 is the indicator that individual *j* is inferred to be individual *i*’s relative.

PC-AiR infers ancestry divergence using 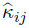 for all pairs of individuals *i, j* ∈ *𝒩* who are not inferred to be relatives. We showed that 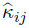 is a consistent estimator of 0 for unrelated pairs with the same ancestry, while unrelated pairs with different ancestry will have 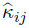 values that are systematically negative. We define a pair of individuals *i* and *j* to be “divergent” if they have different ancestral backgrounds, i.e. 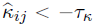, where −*τ_κ_* is the expected lower bound of 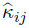 for a pair of unrelated individuals with the same ancestry. Since the distribution of 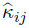 for unrelated pairs with the same ancestry will be symmetric around 0, we expect that the vast majority of these pairs will satisfy 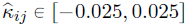 when |*S_ij_*| is large, where 0.025 is the previously mentioned approximate upper bound for unrelated pairs. We have found that setting −*τ_κ_* =−0.025 works well in practice for identifying unrelated pairs of individuals with different admixed ancestries. For all sample individuals *i* ∈ *𝒩*, we calculate 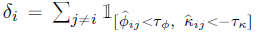, the number of divergent ancestry pairs that individual *i* is a member of. Small *δ_i_* values generally correspond to individuals with ancestry that is similar to the ancestries of many other individuals in *𝒩*, while the highest *δ_i_* values generally correspond to individuals with unique ancestry and/or individuals with an ancestry proportion close to 1 from some subpopulation. Individuals with the highest *δ_i_*values are given priority for inclusion in *U* as they help to ensure that the subset is both diverse and representative of all ancestries in the sample.

The algorithm used by PC-AiR for partitioning the set *𝒩* into subsets *U* and *R* based on measures of ancestry divergence and kinship is presented below:

1. Compute: 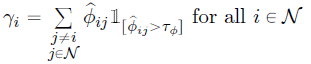.
2. Compute: 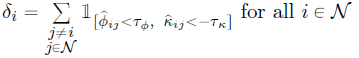.
3. Initialize the two subsets to be *U* = *𝒩* and *R* = Ø, where Ø is the empty set.
4. Compute: 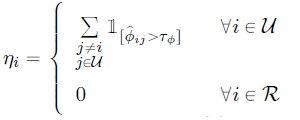. If 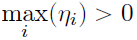, continue to step (5), otherwise go to step (11).
5. Identify 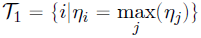, the subset of individuals in *U* with the most relatives in *U*. If |*T*_1_| > 1, where |*T*_1_| is the number of elements in *T*_1_, go to step (6). Otherwise set *T\** = *T*_1_ and go to step (9).
6. Identify 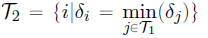, the subset of individuals in *T*_1_ that are members of the least divergent ancestry pairs. If |*T*_2_| > 1, go to step (7). Otherwise set *T\** = *T*_2_ and go to step (9).
7. Identify 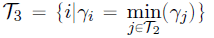, the subset of individuals in *T*_2_ that have the minimum total kinship with their inferred relatives. If |*T*_3_| > 1, go to step (8). Otherwise set *T\** = *T*_3_ and go to step (9).
8. Randomly select one element from *T*_3_ and define this element to be the set *T*\*.
9. Define the sets: *U\** = *U\T*\* and *R*\* = *R*∪*T*\*.
10. Update *U* = *U\** and *R* = *R\** and return to step (4).
11. The algorithm has completed.

This algorithm is both fast and efficient, and the two subsets returned from the algorithm are the ancestry representative and mutually unrelated subset, *U,* and the related subset, *R*, where each individual in *R* has at least one relative in *U*. The algorithm is constructed in such a way that for any subset of mutually related individuals in *𝒩*, one individual in the subset will be chosen to be included in *U,* with priority given to the individual who is a member of the most divergent ancestry pairs. This helps to ensure that every ancestry in *𝒩* is represented by some individual(s) in *U,* while simultaneously satisfying the requirement that individuals in *U* are also mutually unrelated. It also favors selecting individuals with the highest ancestry proportions from each of the *K* subpopulations for *U,* which helps to avoid shrinkage in prediction of principal component values for individuals in *R*, as these individuals will be at the extremes of the *K*−1 dimensional space spanned by the axes of variation representing the ancestries in *𝒩*. Secondary priority for inclusion in *U* is given to individuals that share the most genetic information with their collection of relatives in *𝒩*, also allowing for better prediction of principal component values for relatives in *R*.

### Genetic Similarity Matrix for PC-AiR

The traditional PCA approach for population structure inference with genetic data, e.g., the EIGENSOFT method, performs PCA on standardized genotypes, where the standardized genotype value for individual *i* at SNP *s* is given by

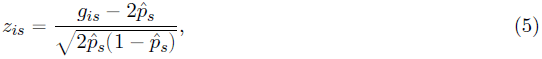

and 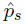 will typically be an allele frequency estimate for SNP *s* calculated using all sample individuals. The PC-AiR method also uses standardized genotypes, but the allele frequencies used for the standardization are calculated using only the unrelated individuals selected for *U*. The standardized genotype values for PC-AiR are calculated from Equation (5) by setting 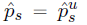, where

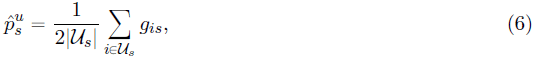

*U_s_* is the subset of individuals in *U* who have non-missing genotype data at SNP *s*, and |*U_s_*| is the number of individuals in *U_s_*. In samples with related individuals and population structure, we have found that using the estimator 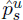 provides better ancestry inference with PC-AiR than using allele frequency estimates calculated from the entire sample, which can be heavily influenced by the correlated genotypes among relatives. For any individual *i* ∈ *𝒩* with a missing genotype value at SNP *s*, *z_is_* is set to 0, i.e. *g_is_* is set equal to 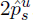, an estimate of its expected value.

Similar minor allele frequency filtering and LD pruning of SNPs that have been recommended for standard PCA (Price et al., 2006; Patterson et al., 2006) should also be used for PC-AiR. Let |*S\**| be the number of SNPs in the pruned and filtered set *S\**, and let *n*, *n_u_*, and *n_r_* be the number of individuals in set *𝒩* and subsets *U* and *R*, respectively, with *n* = *n_u_*+ *n_r_*. We construct **Z**, an *n×*|*S\**| standardized genotype matrix for *𝒩*, with (*i, s*)-th entry equal to *z_is_*, ordered such that the first *n_u_* rows correspond to individuals in *U,* and the remaining *n_r_* rows correspond to individuals in *R*. The standardized genotype matrix for *U* is the *n_u_*×|*S\**| submatrix **Z***_u_* corresponding to the first *n_u_* rows of **Z**. Similarly, the *n_r_*×|*S\**| submatrix **Z***_r_* is the standardized genotype matrix for *R* corresponding to the last *n_r_* rows of **Z**.

Similar to the traditional PCA approach, PC-AiR obtains a genetic similarity matrix (GSM) for population structure inference from standardized genotypes. It is important to note that PCA applied to a GSM that includes all individuals in *𝒩*, as in the traditional PCA approaches, leads to artifactual principal components for ancestry due to confounding from correlated genotypes among relatives, i.e. genetic similarities are reflecting alleles shared IBD among relatives. To protect against confounding caused by sample relatedness, PC-AiR instead calculates a GSM using only the mutually unrelated sample individuals who were selected to be included in the ancestry representative subset, *U*. The empirical *n_u_* × *n_u_* GSM for *U* calculated with the standardized genotype matrix **Z***_u_*is

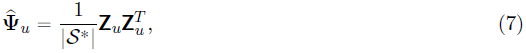

and the (*i, j*)-th entry of 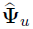 provides a measure of the average genetic similarity across the autosomes for individuals *i, j* ∈ *U*.

### Population Structure Inference in Related Samples with PC-AiR

To obtain principal components that are ancestry representative on a set *𝒩* containing related individuals, the PC-AiR method first performs a PCA using genome-screen data from only those individuals selected to be in the mutually unrelated ancestry representative subset, *U*. PCA is performed by obtaining the eigendecomposition of the GSM 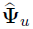 from Equation (7). This procedure sequentially identifies orthogonal axes of variation, i.e. linear combinations of SNPs, that best explain the genotypic variability amongst the individuals in *U,* where each axis of variation reflects the structure that leads to the greatest variability after accounting for the structure explained by all previously defined axes. The eigendecomposition of 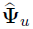 results in an *n_u_ × n_u_* matrix, 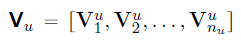, with orthogonal, length *n_u_*, column vectors, and a corresponding length *n_u_* vector, 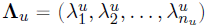, with the property of 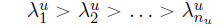. For 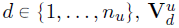 and 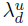 are the corresponding *d^th^* principal component (eigenvector) and eigenvalue of 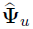, where 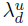 is proportional to the percentage of variability in the genome-screen data for *U* that is explained by 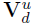. By construction, individuals in *U* are mutually unrelated and have diverse ancestry, so the top principal components of 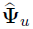 are expected to be representative of ancestry.

Once PCA has been performed on *U,* principal components values for individuals in the related subset, *R*, can be obtained via prediction. Let **L***_u_* be a diagonal matrix created from the vector of eigenvalues, i.e. 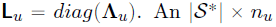 SNP weight matrix giving the relative influence of each SNP on each of the *n_u_* eigenvectors can be obtained as 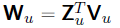, and from the form of the eigendecomposition of the real symmetric matrix 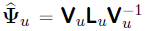, it can be shown (Heath et al., 2008) that the principal components for the ancestry representative subset, *U,* can alternatively be written as:

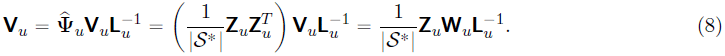

For the related subset, *R*, the PC-AiR method predicts principal components values from Equation (8) by replacing **Z***_u_*, the standardized genotype matrix for individuals in *U,* with **Z***_r_*, the standardized genotype matrix for individuals in *R*. The *n_r_* × *n_u_* matrix of predicted eigenvectors for *R*, which we denote as **Q***_r_*, is thus given by:

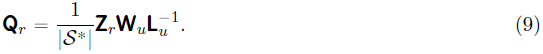

The *d^th^* column in the matrix **Q***_r_*corresponds to PC-AiR’s predicted coordinates along the *d^th^* principal component for the individuals in *R*. We define **Γ** to be the *n* ×*n_u_* matrix of the combined principal components for *U* and *R*, where:

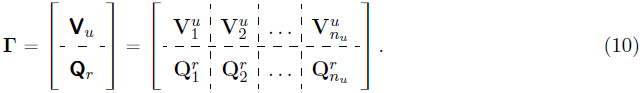

The column vectors of **Γ** are the principal components (axes of variation) of the set *𝒩* = *U* ∪ *R* obtained from the PC-AiR method. The genetic structure that is reflected by all of the principal components for PC-AiR are found using only the ancestry representative subset, *U,* and thus the top principal components from **Γ** are designed to be representative of ancestry in *𝒩*, even in the presence of known or cryptic relatedness.

### Simulation Studies

We perform simulation studies in which both population and pedigree structure are simultaneously present in order to (1) assess the accuracy and robustness of the PC-AiR method for population structure inference in the presence of relatedness, (2) evaluate correction for population stratification with PC-AiR in genetic association studies with cryptic structure, and (3) compare the performance of PC-AiR to existing methods. We simulate a variety of population structure settings, including admixture and ancestry-related assortative mating, with differentiation between populations ranging from subtle to large. We evaluate population structure inference for four different relationship configurations, where each configuration corresponds to a specific setting of genealogical relationships among the sample individuals. In all simulation studies considered, pedigree information on the sample individuals is hidden and genetic relatedness is inferred from the genotype data with the PC-AiR method using the KING-robust kinship estimator in Equation (1).

### Population Structure Settings

The population structure settings we consider are similar to the settings in Price et al. (2006), where PCA was performed with the EIGENSOFT software in unrelated samples for inference on and adjustment for population structure in GWAS, except that our simulation studies include related individuals. We consider population structure settings where individuals have ancestry derived from two populations, and the allele frequencies at 100,000 SNPs for each of these two populations are generated using the Balding-Nichols model (Balding and Nichols, 1995). More precisely, for each SNP *s*, the allele frequency *p_s_* in the ancestral population is drawn from a uniform distribution on [0.1, 0.9], and the allele frequency in population *k* ∈ {1, 2} is drawn from a beta distribution with parameters 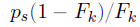 and 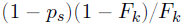, where the quantity *F_k_* is equivalent to Wright’s measure of population differentiation (Wright, 1949) from the ancestral population. In all simulations, we set *F*_1_ and *F*_2_ equal to a common value, *F_ST_*. To generate allele frequencies derived from populations ranging from closely related to highly divergent, we consider *F_ST_* values from 0.01 to 0.2.

For each *F_ST_* value considered, we simulate three population structure settings. Population structures I and II both consist of individuals sampled from an admixed population formed from populations 1 and 2. For population structure I, all unrelated individuals and pedigree founders have ancestry proportions *a* from population 1 and (1−*a*) from population 2, with the parameter *a* for each individual drawn from a uniform distribution on [0, 1]. Population structure II is similar to population structure I, but with the ancestry parameter, *a*, drawn from a beta distribution with mean 0.4 and standard deviation 0.1 for 50% of the unrelated individuals and pedigree founders, and with mean 0.6 and standard deviation 0.1 for the other 50%. All founders within the same pedigree have *a* drawn from the same beta distribution for population structure II. Population structure III consists of non-admixed individuals, where 50% of the unrelated individuals and pedigrees are sampled from population 1, and the other 50% are sampled from population 2. Both population structure settings II and III have ancestry-related assortative mating, i.e., the mating of founder individuals in every pedigree occurs with individuals who have either the same (population structure III) or similar (population structure II) ancestry, while population structure I has random mating that is independent of ancestry.

### Relationship Configurations

Three of the four relationship configurations simulated include both related and unrelated individuals. Relationship configuration I consists of 200 unrelated individuals and 200 individuals from 10 four-generation pedigrees, where each pedigree has a total of 20 individuals (Figure S1). Relationship configuration II is comprised of 280 unrelated individuals with 20 parent-offspring trios, and relationship configuration III includes 260 unrelated individuals with 20 sibling pairs. To sample pedigree relationships within a given setting of population structure, we simulate genotypes for pedigree founders under Hardy-Weinberg equilibrium (HWE) according to the chosen population structure setting and then drop alleles down the pedigree. Relationship configuration IV consists of 320 unrelated individuals without any family structure. We include the unrelated sample setting in our simulation studies in order to evaluate any loss in population structure inference with the PC-AiR method compared to standard PCA in a setting where standard PCA is appropriate and has been previously demonstrated to perform well.

## Results

### Subtle Population and Pedigree Structure

We first considered samples with subtle population structure, where the ancestry of the sample individuals was derived from two closely related populations. We set *F_ST_* to 0.01 (a typical value for divergent European populations) and generated genotype data under population structure I for each of the four relationship configurations. Population structure inference with PC-AiR was compared to that of standard PCA with the EIGENSOFT software. To assess the performance of the two methods, we included the top principal components (axes of variation) from each method as predictors for the true simulated ancestry of the sample individuals in a linear regression model, and the proportion of ancestry explained, as measured by *R*^2^, was used to evaluate prediction accuracy. We also compared the efficiency of PC-AiR to EIGENSOFT by assessing the number of top axes of variation required to attain an *R*^2^ of at least 0.99 for ancestry. It should be noted that since the data in the simulation studies contained only one added dimension of population structure, an optimal method would require only a single axis of variation for complete ancestry inference. Both PC-AiR and EIGENSOFT were provided only genotype data without any additional pedigree information on the sample individuals.

Figure 1 displays the population structure inference results for relationship configuration I from both PC-AiR and EIGENSOFT. Figure 1B displays the top two axes of variation obtained by EIGENSOFT, which almost entirely reflected pedigree structure in the sample. The ten spikes of points radiating from the center cluster in the figure correspond to the individuals who are members of the ten pedigrees, and the cluster of points in the center of the plot corresponds to the 200 individuals who do not have any relatives in the sample. In contrast, the top two axes of variation from PC-AiR were not confounded by family structure, as illustrated in Figure 1A, and the top axis explained ancestry in the sample nearly perfectly, with an *R*^2^ of 0.993 (Figure 1C). Figure 1D shows that the top axis of variation from EIGENSOFT did not reflect population structure and did not adequately capture the ancestry of the sample individuals, with an *R*^2^ of only 0.133. The efficiency for population structure inference of both methods is illustrated in Figure 1E, where the proportion of ancestry explained (*R*^2^ values) for each of the top axes of variation is displayed. EIGENSOFT required the top 51 axes to be included as predictors in a linear regression model to achieve an *R*^2^ of at least 0.99 for ancestry. In contrast, a single axis of variation from PC-AiR had an *R*^2^ greater than 0.99, thus demonstrating a substantial improvement in efficiency with PC-AiR over EIGENSOFT in this setting with both subtle population structure and relatedness.

**Figure 1.**
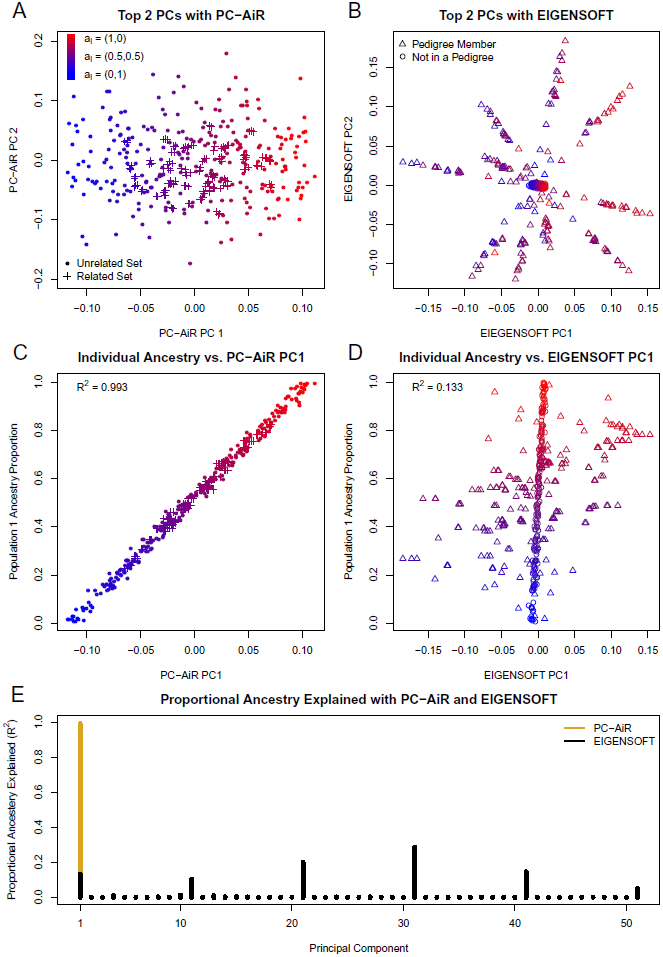
Comparison of PC-AiR and EIGENSOFT for Data Simulated under Relationship Configuration I and Population Structure I with *F_ST_* = 0.01. (A and B) Scatter plots of principal components 1 and 2 from PC-AiR (A) and EIGENSOFT (B), respectively. (C and D) Scatter plots of the simulated population 1 ancestry proportions vs. coordinates along principal component 1 for each individual from PC-AiR (C) and EIGENSOFT (D), respectively. (A-D) The color of each point represents that individual’s true ancestry; red for population 1, blue for population 2, and an intermediate color for an admixed individual. (A and C) A dot represents an individual in the mutually unrelated ancestry representative set, and a plus represents an individual in the related set. (B and D) A circle represents an individual not in a pedigree, and a triangle represents an individual who is a member of a pedigree. (E) Barplot of the efficiency of PC-AiR and EIGENSOFT. Each bar represents the proportion of ancestry explained (*R*^2^ value) by each principal component from PC-AiR (gold) and EIGENSOFT (black), until a cumulative *R*^2^ of 0.99 is achieved.

Population structure inference results with PC-AiR and EIGENSOFT for relationship configurations II and III are presented in Figures S2 and S3. The top axes of variation from EIGENSOFT were influenced by relatedness, as expected; however, since relationship configurations II and III have substantially less pedigree structure than relationship configuration I, there was some improvement in ancestry prediction with the top axis in each of these two settings, with *R*^2^ values of 0.870 and 0.933, respectively. For both relationship configurations II and III, the top 21 axes of variation from EIGENSOFT were required to attain an *R*^2^ of at least 0.99 for predicting ancestry. In comparison, the PC-AiR analysis was robust to the relatedness in the sample, and the single top axis of variation for both relationship configurations II and III attained an *R*^2^ value greater than 0.99 for predicting ancestry. For relationship configuration IV, PC-AiR accurately identified all sample individuals to be unrelated, i.e. the ancestry informative subset, *U,* was the entire sample, *𝒩*, so the PC-AiR method reduced to standard PCA, and inference with either PC-AiR or EIGENSOFT was essentially identical. The *R*^2^ for ancestry with the top axis of variation from both methods was greater than 0.99, illustrating that there is no loss in accuracy or efficiency compared to standard PCA when using PC-AiR for population structure inference in samples where all individuals are unrelated.

We also evaluated the performance of PC-AiR and EIGENSOFT under population structures II and III with *F_ST_* set to 0.01 for each of the relationship configurations. The results are given in Table 1, and the conclusions drawn from these population structure settings are the same as those for population structure I. For the three relationship configurations that included related samples, a single axis of variation from PC-AiR fully explained the ancestry in the sample and provided better prediction of ancestry than using ten (or more) axes from EIGENSOFT. For relationship configuration IV, where all sample individuals were unrelated, PC-AiR and EIGENSOFT gave essentially identical results, with the top axis from both methods fully explaining the true ancestry.

**Table 1:** Proportion of Ancestry Explained (*R*^2^) by PC-AiR and EIGEN-SOFT in Simulation Studies

| Relationship Configuration | Population Structure | $F_{ST}$ | PC-AiR | EIGENSOFT | | | |
| --- | --- | --- | --- | --- | --- | --- | --- |
| | | | $d^a = 1$ | $d = 1$ | $d = 4$ | $d = 10$ | $d^{*b}$ |
| I | I | 0.01 | 0.993 | 0.133 | 0.145 | 0.165 | 51 |
|  |  | 0.1 | 0.999 | 0.977 | 0.977 | 0.993 | 10 |
|  |  | 0.2 | 0.999 | 0.995 | 0.995 | 0.999 | 1 |
|  | II | 0.01 | 0.949 | 0.302 | 0.360 | 0.402 | 51 |
|  |  | 0.1 | 0.998 | 0.741 | 0.741 | 0.755 | 41 |
|  |  | 0.2 | 0.999 | 0.882 | 0.882 | 0.914 | 21 |
|  | III | 0.01 | 0.999 | 0.832 | 0.832 | 0.832 | 22 |
|  |  | 0.1 | 0.999 | 0.998 | 0.998 | 0.998 | 1 |
|  |  | 0.2 | 0.999 | 0.999 | 0.999 | 0.999 | 1 |
| II | I | 0.01 | 0.994 | 0.870 | 0.871 | 0.872 | 21 |
|  |  | 0.1 | 0.999 | 0.999 | 0.999 | 0.999 | 1 |
|  |  | 0.2 | 0.999 | 0.999 | 0.999 | 0.999 | 1 |
|  | II | 0.01 | 0.942 | 0.259 | 0.313 | 0.320 | 21 |
|  |  | 0.1 | 0.998 | 0.983 | 0.983 | 0.984 | 21 |
|  |  | 0.2 | 0.999 | 0.996 | 0.996 | 0.996 | 1 |
|  | III | 0.01 | 0.999 | 0.990 | 0.990 | 0.990 | 1 |
|  |  | 0.1 | 0.999 | 0.999 | 0.999 | 0.999 | 1 |
|  |  | 0.2 | 0.999 | 0.999 | 0.999 | 0.999 | 1 |
| III | I | 0.01 | 0.990 | 0.933 | 0.933 | 0.936 | 21 |
|  |  | 0.1 | 0.999 | 0.999 | 0.999 | 0.999 | 1 |
|  |  | 0.2 | 0.999 | 0.999 | 0.999 | 0.999 | 1 |
|  | II | 0.01 | 0.922 | 0.220 | 0.230 | 0.250 | 21 |
|  |  | 0.1 | 0.997 | 0.992 | 0.992 | 0.993 | 1 |
|  |  | 0.2 | 0.999 | 0.998 | 0.998 | 0.998 | 1 |
|  | III | 0.01 | 0.998 | 0.995 | 0.995 | 0.995 | 1 |
|  |  | 0.1 | 0.999 | 0.999 | 0.999 | 0.999 | 1 |
|  |  | 0.2 | 0.999 | 0.999 | 0.999 | 0.999 | 1 |

<sup>a</sup> $d$ denotes the number of axes of variation included as predictors in the linear regression model to determine the $R^2$ value for either PC-AiR or EIGENSOFT.
<sup>b</sup> $d^*$ is the number of axes of variation from EIGENSOFT that are required to either match the $R^2$ value of the first axis of variation from PC-AiR or achieve an $R^2$ of 0.99, whichever is smaller.

### Relatedness and Admixture from Divergent Populations

We also conducted simulation studies with relatedness and admixture from divergent populations. We considered relationship configuration I and population structure II, where we set *F_ST_* to 0.1 (a value representative of continental-level ancestry differences) in the Balding-Nichols model to simulate allele frequencies at SNPs derived from two divergent populations. We evaluated and compared the performance of PC-AiR to PCA with the EIGENSOFT software, MDS with the PLINK software, and the two model-based methods ADMIXTURE and FRAPPE for proportional ancestry estimation. As in the previous subsection, no genealogical information on the sample individuals was provided to any of the analysis methods, so the FamPCA method could not be used as it is restricted to settings with known pedigrees. The ADMIXTURE and FRAPPE software analyses were conducted with the correct number of populations specified.

The population structure inference results for each method considered are shown in Figure 2, where each panel is a plot of the simulated population 1 ancestry proportions against the inferred ancestry from one of the methods. The top axis of variation from PC-AiR had an *R*^2^ of 0.998 and provided nearly perfect inference on ancestry for the sample individuals (Figure 2A). Similar to the EIGENSOFT results for the simulations with subtle population structure and relatedness, the top axis of variation did not adequately reflect the ancestry in this related sample with admixture from divergent populations, attaining an *R*^2^ of only 0.741 (Figure 2B). ADMIXTURE and FRAPPE gave identical ancestry proportion estimates for all individuals in the simulation, and Figure 2D shows estimated proportional ancestry plotted against the simulated ancestry proportions from population 1. These model-based ancestry estimation methods were confounded by the pedigree structure in the sample and performed similarly to PCA, with an *R*^2^ of only 0.730. While the top dimension of MDS achieved an *R*^2^ of 0.785 and provided some improvement in predicting ancestry over both of the model-based methods as well as the top axis of variation from EIGENSOFT, it was also confounded by sample relatedness, as shown in Figure 2C.

**Figure 2.**
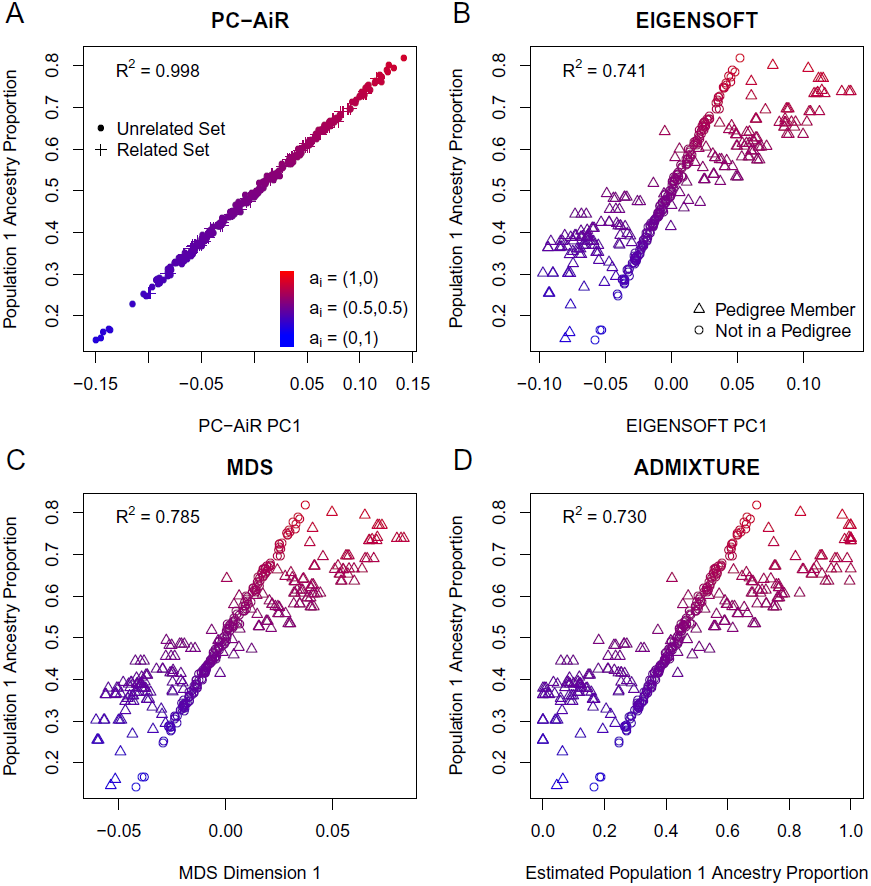
Comparison of Population Structure Inference Results for Data Simulated under Relationship Configuration I and Population Structure II with *F_ST_* = 0.1. Scatter plots of the simulated population 1 ancestry proportions for each individual are plotted against: (A) coordinates along principal component 1 from PC-AiR, (B) coordinates along principal component 1 from EIGENSOFT, (C) coordinates along dimension 1 from MDS, and (D) the estimated ancestry proportions from ADMIXTURE for the inferred population with the highest *R*^2^. The color of each point represents that individual’s true ancestry; red for population 1, blue for population 2, and an intermediate color for an admixed individual. (A) A dot represents an individual in the mutually unrelated ancestry representative set, and a plus represents an individual in the related set. (B-D) A circle represents an individual not in a pedigree, and a triangle represents an individual who is a member of a pedigree.

We also evaluated the performance of PC-AiR and EIGENSOFT for all combinations of relationship configurations and population structure settings with *F_ST_* set to 0.1 and 0.2 (Table 1). For all settings considered, the top axis of variation from PC-AiR gave nearly perfect ancestry inference, attaining an *R*^2^ > 0.99. The extent to which EIGENSOFT’s PCA was confounded by the relatedness depended on how divergent the populations were, i.e. the *F_ST_* values, and how complex the pedigree structure was; however, a single axis of variation from PC-AiR always performed as well as or better than using ten axes of variation from EIGENSOFT for ancestry prediction.

### Ancestry Inference in Related Samples with Reference Panels

Reference population panels are commonly used for improved ancestry inference in unrelated samples from admixed populations, such as African Americans and Hispanics. We conducted a simulation study evaluating population structure inference with reference panels in admixed samples with relatedness. We considered the same simulation study discussed in detail in the previous subsection, but we now included reference panels consisting of 50 unrelated individuals randomly sampled from each of the two populations. The same population structure methods from the previous subsection were used, and the results are displayed in Figure S4. Ancestry inference with EIGENSOFT, MDS, ADMIXTURE, and FRAPPE was substantially improved by including the reference panels as compared to the analyses without them, but PC-AiR still outperformed all methods, with the top axis of variation achieving an *R*^2^ of 0.999 with ancestry. The supervised analyses with ADMIXTURE and FRAPPE including the reference panels gave identical results to each other, as in the unsupervised analyses without reference panels, and the estimated ancestry proportions had an *R*^2^ of 0.973 with the simulated ancestries. Similarly, the top axis of variation from each of EIGENSOFT and MDS reached *R*^2^ values of 0.970 and 0.979 respectively.

Interestingly, the top axis of variation from PC-AiR without additional reference population samples had an *R*^2^ of 0.998 and provided better ancestry prediction than all of the competing methods with the reference panels. Even with the inclusion of reference panels, there remains some bias in ancestry inference for all methods, except for PC-AiR, that is induced by the presence of related individuals in the sample. This can be seen in Figure S4, where the inferred ancestries for individuals with relatives in the sample were systematically biased for each of the competing methods. We have found that using ADMIXTURE (or FRAPPE) to conduct separate individual ancestry analyses for each of the admixed sample individuals with the reference panels can remove the bias caused by sample relatedness, known or cryptic, as long as the reference panel samples are appropriate surrogates for the underlying populations. We performed separate individual ancestry analyses with ADMIXTURE for each sample individual, where each analysis included genotype data from a single admixed sample individual and all individuals in the two reference population panels, and the estimated ancestries attained an *R*^2^ of 0.999, the same as PC-AiR.

### Correcting for Structure in Genetic Association Studies

We also performed simulation studies to compare population structure correction in genetic association studies with PC-AiR to existing approaches. Allele frequencies were generated at 100,000 null SNPs for two ancestral populations with *F_ST_* set to 0.1. We define 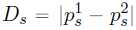 to be the absolute difference in the reference allele frequencies between ancestral populations 1 and 2 at SNP *s*. We also define three classes of SNPs based on *D_s_*, where SNPs with *D_s_ <* 0.2, 0.2 ⩽ *D_s_* < .4, and *D_s_* ⩾ 0.4 were considered to have “low differentiation,” “moderate differentiation,” and “high differentiation,” respectively. Of these 100,000 SNPs, approximately 70% had low differentiation, 25% were moderate, and 5% were highly differentiated. Genotype data was generated under population structure II for sample individuals related according to relationship configuration I, and for each individual *i* in the sample, a quantitative trait value *y_i_* was simulated according to the model 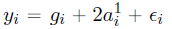, where 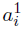 is the genome-wide ancestry proportion from population 1 for individual *i*, *g_i_* is the number of alleles individual *i* has at the causal SNP, and ∈*_i_ ∼* N(0, 1) is a random environmental effect assumed to be acting independently on individuals. The frequency of the selected casual variant in populations 1 and 2 was 0.13 and 0.17, respectively.

The following statistical methods were evaluated for genetic association testing: (1) linear regression without ancestry adjustment, (2) EIGENSTRAT, (3) linear regression with principal components from PC-AiR included as fixed effects, (4) GEMMA (Zhou and Stephens, 2012) and EMMAX (Kang et al., 2010), which are “exact” and “approximate” linear mixed effects model methods, respectively, that use an empirical genetic relatedness matrix to capture both population and pedigree sample structure, and (5) EMMAX with principal components from EIGENSOFT or PC-AiR included as fixed effects. For the association analyses, each null SNP was included as a fixed effect in the statistical models and was tested for association with the simulated quantitative trait. The genomic control inflation factor (Devlin and Roeder, 1999) *λ_GC_* was used to evaluate confounding due to unaccounted for sample structure, where *λ_GC_* ≈ 1 indicates appropriate correction for population and family structure, while *λ_GC_* > 1 indicates elevated type-I error.

The results of the simulations are given in Table 2. As expected, all of the association tests using linear regression models have inflated type 1 error since these methods either (1) do not account for any of the sample structure, or (2) account for population structure but not relatedness. Including a single principal component from PC-AiR in the linear regression model results in a lower *λ_GC_* compared to EIGENSTRAT with the top ten principal components for all classes of SNPs. This is because the top PC from PC-AiR nearly perfectly explains ancestry (*R*^2^ = 0.998), while the top 10 PCs from EIGENSOFT have an *R*^2^ of only 0.672 for ancestry as a result of the relatedness in the sample. The mixed model approaches considered, EMMAX and GEMMA, are also not properly calibrated, with *λ_GC_* > 1 for SNPs with moderate to high differentiation, and *λ_GC_* < 1 for SNPs with low differentiation. EMMAX with the top ten PCs from EIGENSOFT included as fixed effects is still not properly calibrated due to incomplete correction for population stratification. However, including a single PC from PC-AiR as a fixed effect with EMMAX results in appropriate calibration of the association test statistics, with *λ_GC_* = 1 for all classes of SNPs.

**Table 2:** Genomic Control *λ_GC_* for Association Testing Simulation Study

| Method | Highly<sup>a</sup><br>Differentiated | Moderately<sup>b</sup><br>Differentiated | Lowly<sup>c</sup><br>Differentiated |
| --- | --- | --- | --- |
| Linear Regression | 3.16 | 1.82 | 1.25 |
| EIGENSTRAT with 10 PCs | 1.19 | 1.12 | 1.07 |
| Linear Reg. + 1 PC from PC-AiR | 1.04 | 1.05 | 1.05 |
| GEMMA | 1.35 | 1.10 | 0.95 |
| EMMAX | 1.32 | 1.08 | 0.94 |
| EMMAX + 10 PCs from EIGENSOFT | 1.07 | 1.02 | 0.98 |
| EMMAX + 1 PC from PC-AiR | 1.00 | 1.00 | 1.00 |

<sup>a</sup> Highly differentiated SNPs have allele frequency differences $\geq 0.4$ between the two ancestral populations.
<sup>b</sup> Moderately differentiated SNPs have allele frequency differences $< 0.4$ and $\geq 0.2$ between the two ancestral populations.
<sup>c</sup> Lowly differentiated SNPs have allele frequency differences $< 0.2$ between the two ancestral populations.

### Population Structure Inference in Admixed HapMap Samples

#### HapMap MXL Data

We analyzed high-density genotype data from the Mexican Americans in Los Angeles, California (MXL) population sample of HapMap 3 for population structure inference. We applied PC-AiR, EIGENSOFT, MDS, ADMIXTURE, and FamPCA to the 86 genotyped individuals, and we compared the population structure inference results of these methods to a supervised individual ancestry estimation analysis with ADMIXTURE that included continental reference population panels. For the supervised analysis with ADMIXTURE, the number of ancestral populations was set to 3, for which the HapMap CEU (Utah residents with ancestry from northern and western Europe from the Centre d’Etude du Polymorphisme Human collection) and YRI (Yoruba in Ibadan, Nigeria) samples were included as the reference population panels for European and African ancestry, respectively, and for which the Human Genome Diversity Project (HGDP) (Li et al., 2008) samples from the Americas were included for Native American ancestry. The analyses were based on 150,872 autosomal SNPs that were genotyped in both the HapMap and HGDP datasets. To protect against potential confounding due to relatedness in the supervised ancestry analysis, a separate ADMIXTURE analysis was conducted for each of the HapMap MXL individuals, where each analysis included a single HapMap MXL individual and the reference population panels. All methods, except for FamPCA, were only provided the SNP genotype data on the sample individuals for population structure inference, without any additional information on the pedigree relationships. The FamPCA method was also provided the documented pedigrees in the HapMap MXL which includes 24 genotyped trios, 5 families with two genotyped individuals, and 4 families with a single genotyped individual. The PC-AiR method used the KING-robust kinship coefficient estimator in Equation (1) and the relatedness threshold *τ_φ_* = 0.025 to infer genetic relatedness in the sample, and a MAF filter of 5% was used on SNPs for population structure inference.

Figure 3F presents a bar plot of the results from the supervised individual ADMIXTURE ancestry analysis. In the bar plot of ancestry proportion estimates, individuals (vertical bars) are arranged in increasing order (left to right) of genome-wide European ancestry proportion. Our proportional ancestry estimates were similar to the results from a previous supervised analysis of this data (Thornton et al., 2012; Gravel et al., 2013). HapMap MXL individuals have modest African ancestry with little variation, with a mean of 6% and a standard deviation (SD) of 1.8%. The sample individuals are largely derived from European and Native American ancestry, with means of 49.9% (SD = 14.8%) and 44.1% (SD = 14.8%) respectively. Since the European and Native American ancestry proportions are predominant, nearly perfectly negatively correlated (with a correlation of −0.99), and quite variable, ranging from 18.0% to 91.0% and from 4.2% to 80.4% respectively, we expected that an optimal population structure inference method would require only a single axis of variation to explain these two ancestries in the HapMap MXL.

**Figure 3.**
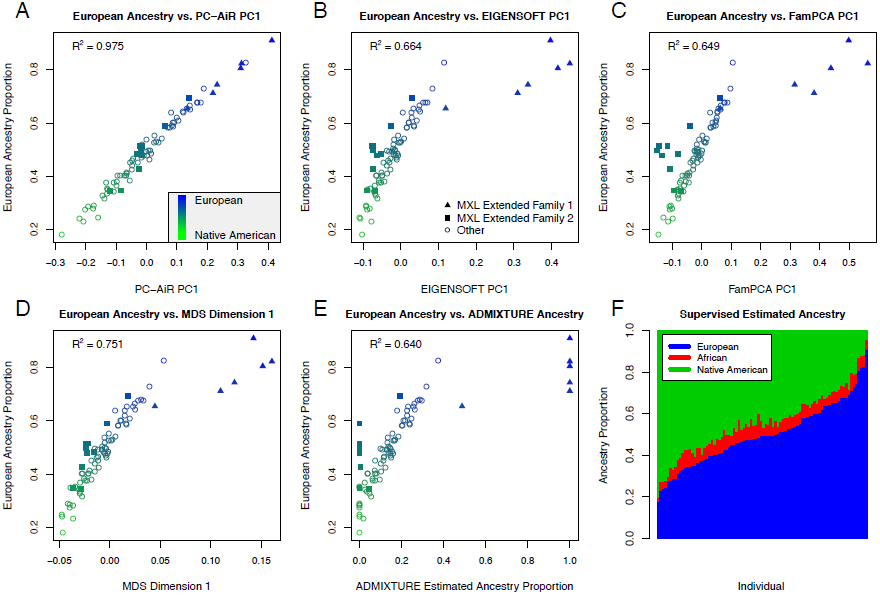
Comparison of Population Structure Inference for the HapMap MXL Sample. (F) Individual ancestry estimates for 86 HapMap MXL samples from a supervised individual ancestry analysis with ADMIXTURE. Each individual is represented by a vertical bar; estimated European (HapMap CEU), African (HapMap YRI), and Native American (HGDP samples from the Americas) ancestry proportions are shown in blue, red, and green, respectively. Scatter plots of the European ancestry proportions estimated from a supervised individual ancestry analysis with ADMIXTURE for each individual are plotted against: (A) coordinates along principal component 1 from PC-AiR, (B) coordinates along principal component 1 from EIGENSOFT, (C) coordinates along principal component 1 from FamPCA, (D) coordinates along dimension 1 from MDS, and (E) the estimated ancestry proportions from an unsupervised analysis with ADMIXTURE for the inferred population with the highest *R*^2^. The color of each point represents that individual’s ancestry as estimated from the supervised individual ancestry analysis with ADMIXTURE; blue for European, green for Native American, and an intermediate color for an admixed individual. Individuals who are members of MXL Extended Family 1 or 2 are plotted as triangles or squares, respectively, and remaining individuals are plotted as circles.

The population structure inference results for European and Native American ancestry in the HapMap MXL are given in Table 3. PC-AiR’s top axis of variation was nearly perfectly correlated with European (and Native American) ancestry, as estimated from the supervised individual ADMIXTURE ancestry analysis, with an *R*^2^ of 0.98 (Figure 3A). In contrast, the top axis of variation from each of EIGENSOFT, FamPCA, and MDS had an *R*^2^ for European ancestry of only 0.66, 0.65, and 0.75 respectively. For the unsupervised ADMIXTURE analysis that did not include reference panels, the highest *R*^2^ for either European or Native American ancestry with any estimated ancestry component was only 0.64. Figures 3B, 3C, 3D, and 3E illustrate that ancestry inference in the HapMap MXL for each of these competing methods was confounded by relatedness, including the FamPCA method, which was provided the documented pedigree relationships. Ancestry inference with FamPCA was confounded by cryptic relatedness present in the HapMap MXL including a previously reported (Thornton et al., 2012) extended pedigree consisting of two smaller documented pedigrees, which we have labeled in Figure 3 as MXL Extended Family 1. Without being provided any pedigree information, a single axis of variation from PC-AiR gave better prediction of both European and Native American ancestry than the top ten axes from EIGENSOFT, MDS, and FamPCA, as shown in Table 3. Remarkably, the top axis of variation from PC-AiR without using any reference population samples gave comparable ancestry inference on European and Native American ancestry to a supervised ancestry analysis that included reference panels, similar to the results from the simulation studies.

**Table 3:** Population Structure Inference Results for HapMap MXL and ASW

| $R^2$ Values | | | | | | |
| --- | --- | --- | --- | --- | --- | --- |
| Ancestry | $d^a$ | PC-AiR | EIGENSOFT<sup>b</sup> | MDS<sup>c</sup> | FamPCA<sup>d</sup> | ADMIXTURE<sup>e</sup> |
| MXL Sample |  |  |  |  |  |  |
| European | 1 | 0.975 | 0.664 | 0.751 | 0.649 | 0.640 |
|  | 4 | - | 0.914 | 0.935 | 0.969 | - |
|  | 10 | - | 0.924 | 0.943 | 0.970 | - |
| Native American | 1 | 0.977 | 0.661 | 0.748 | 0.651 | 0.633 |
|  | 4 | - | 0.908 | 0.929 | 0.968 | - |
|  | 10 | - | 0.911 | 0.932 | 0.969 | - |
| MXL + ASW Sample |  |  |  |  |  |  |
| European | 2 | 0.988 | 0.858 | 0.892 | 0.862 | 0.615 |
|  | 4 | - | 0.868 | 0.899 | 0.878 | - |
|  | 10 | - | 0.963 | 0.970 | 0.987 | - |
| Native American | 2 | 0.995 | 0.953 | 0.962 | 0.951 | 0.866 |
|  | 4 | - | 0.958 | 0.967 | 0.961 | - |
|  | 10 | - | 0.987 | 0.989 | 0.996 | - |
| African | 2 | 0.999 | 0.996 | 0.997 | 0.997 | 0.990 |
|  | 4 | - | 0.996 | 0.997 | 0.997 | - |
|  | 10 | - | 0.999 | 0.999 | 0.999 | - |

Population structure inference results from each method were compared to ancestry estimates from a supervised individual ancestry analysis with ADMIXTURE including reference population panels.
<sup>a</sup> $d$ denotes the number of axes of variation included as predictors in the linear regression model to determine the $R^2$ value for each of the methods.
<sup>b</sup> PCA was performed with the EIGENSOFT software.
<sup>c</sup> MDS was implemented in the PLINK software.
<sup>d</sup> FamPCA is the Zhu et al. (2008) method as implemented in the KING (Manichaikul et al., 2010) software.
<sup>e</sup> An unsupervised ADMIXTURE analysis was conducted without including reference population panels. FRAPPE results were identical to ADMIXTURE.

#### Combined HapMap ASW and MXL Data

To evaluate the performance of the population structure inference methods in an admixed population structure setting with three predominant continental ancestries and relatedness, we considered an analysis of the combined HapMap ASW (African American individuals in the southwestern USA) and MXL samples. Similar to our ancestry estimation analysis of the HapMap MXL, we also conducted a supervised individual ADMIXTURE analysis for the 87 genotyped individuals in the HapMap ASW with reference population panels included for European, Native American, and African ancestries. Figure S5A shows a barplot of the results from the supervised individual ADMIXTURE ancestry analysis of the HapMap MXL and ASW samples, which illustrates that these populations have very different ancestral backgrounds. Most of the HapMap ASW ancestry is African, with a mean of 77.5% (SD = 8.4%). There is also a large European ancestry component, with a mean of 20.5% (SD = 7.9%); however, unlike the HapMap MXL, there is very little Native American ancestry in the HapMap ASW, with a mean of only 1.9% (SD = 3.5%). Since there are three predominant continental ancestries in the combined HapMap ASW and MXL samples, we expected that an optimal method would require two axes of variation to fully explain continental population structure.

We applied each of the dimension reduction methods (i.e. PC-AiR, EIGENSOFT, MDS, and FamPCA) to the combined HapMap ASW and MXL samples and compared the results to the supervised individual ancestry analysis with ADMIXTURE that included the reference population panels; results are shown in Table 3. All of the methods were able to fully explain the African ancestry with two axes of variation, achieving *R*^2^ values greater than 0.99. For European ancestry, PC-AiR’s top two axes of variation achieved an *R*^2^ value of 0.99, while the top two axes from each of the competing population structure methods had *R*^2^ values less than 0.90. With an *R*^2^ value greater than 0.99, PC-AiR’s top two axes of variation also explained Native American ancestry better than the top two axes from EIGENSOFT, MDS, and FamPCA, with corresponding *R*^2^ values of 0.95, 0.96, and 0.95, respectively. These results are illustrated in Figure 4, where we can see that the top two axes of variation from each of these methods, except PC-AiR, were confounded by relatedness. In fact, the top ten axes of variation from EIGENSOFT, MDS, and FamPCA were highly confounded by pedigree structure, whereas axes beyond the top two from PC-AiR did not represent any identifiable structure and appear to be noise (Figures S6 - S10). As a consequence, the top ten axes of variation from both EIGENSOFT and MDS were not able to explain European and Native American ancestry as well as the top two axes from PC-AiR. Intersetingly, FamPCA required ten axes of variation to match PC-AiR’s top two, despite FamPCA being provided the documented pedigree information for both the HapMap MXL and ASW samples (Table 3). PC-AiR appropriately accounted for both the known and cryptic relatedness in the sample for optimal and efficient inference on ancestry with only two axes of variation.

**Figure 4.**
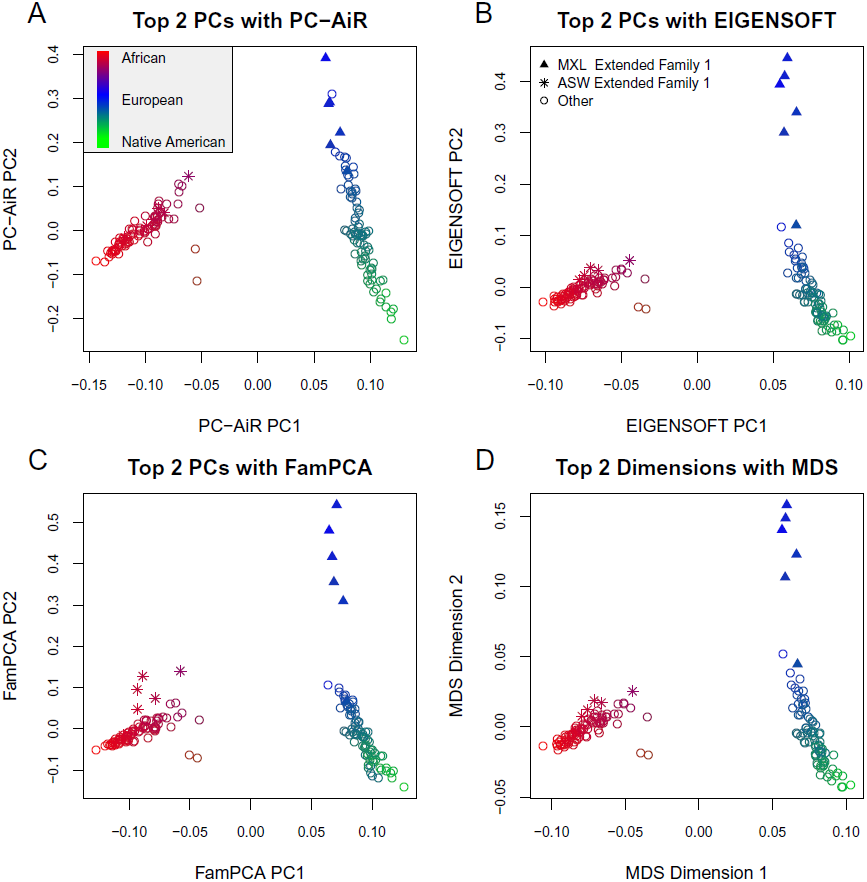
Comparison of Population Structure Inference for the HapMap MXL and ASW Combined Sample. Scatter plots of the top two axes of variation from PC-AiR (A), EIGENSOFT (B), FamPCA (C), and MDS (D). The color of each point represents that individual’s ancestry as estimated from a supervised individual ancestry analysis with ADMIXTURE; blue for European (HapMap CEU), red for African (HapMap YRI), green for Native American (HGDP samples from the Americas), and an intermediated color for an admixed individual. Individuals who are members of MXL Extended Family 1 or ASW Extended Family 1 are plotted as triangles or stars, respectively, and remaining individuals are plotted as circles.

We also performed an unsupervised ancestry analysis with ADMIXTURE and FRAPPE without including reference panel samples and we compared the results to the supervised ADMIXTURE analysis. ADMIXTURE and FRAPPE performed identically to each other, as expected, and a barplot of the estimated ancestry proportions from the unsupervised ancestry analysis is given in Figure S5B. Two of the three components of ancestry essentially distinguish the ASW from the MXL samples, while the third was completely confounded by pedigree structure. This estimated ancestry components were able to attain an *R*^2^ value of 0.99 for African ancestry, but the *R*^2^ values were only 0.87 for Native American ancestry and 0.62 for European ancestry, thus performing the worst of all the methods for ancestry inference in the combined HapMap MXL and ASW samples.

### Assessment of Computation Time

The computation time for PC-AiR depends on both the sample size and the number of markers being analyzed. To analyze a simulated sample of 800 individuals, where 400 individuals are from 20 pedigrees and the remaining 400 individuals are unrelated, with 100 K, 50 K, and 20 K SNPs required 28.5s, 14.8s, and 6.3s, respectively, on a 2.5 GHz laptop with 8 GB memory. The PC-AiR analysis of the HapMap data with 150,872 SNPs required 1.8s for the MXL sample with 86 individuals and 3.9s for the combined ASW and MXL sample with 173 individuals. All computation times refer to the time to run the PC-AiR algorithm, and do not include the time to estimate the measure of relatedness and divergence. The KING software implements a highly efficient algorithm for obtaining relatedness/divergence estimates, and evaluating millions of pairs of individuals in a sample can be conducted in a matter of minutes.

## Discussion

Genetic ancestry inference has been motivated by a variety of applications in population genetics, genetic association studies, and other genomic research areas. Advancements in array-based genotyping technologies have largely facilitated the investigation of genetic diversity at remarkably high levels of detail, and a variety of methods have been proposed for the identification of genetic ancestry differences among unrelated sample individuals using high-density genome-screen data. It is common, however, for genetic studies to have sample structure that is due to both population stratification and relatedness, and existing population structure inference methods can fail in related samples. We develop PC-AiR, a method for robust population structure inference in the presence of known or cryptic relatedness. PC-AiR applies a computationally efficient algorithm that uses pairwise measures of kinship and ancestry divergence from genome-screen data for the identification of a diverse subset of mutually unrelated individuals that is representative of the ancestries in the entire sample. Principal components that are representative of ancestry are obtained by performing PCA directly on genotype data from individuals selected for the ancestry representative subset, while coordinates along the axes of variation for the remaining individuals in the sample are predicted based on genetic similarities with the diverse subset. The PC-AiR method does not require the genealogy of the sampled individuals to be known, and it can be used across a variety of study designs, ranging from population based studies where individuals are assumed to be unrelated to family based studies with partially or completely unknown pedigrees.

In simulation studies with a broad range of population structure settings, including ancestry admixture, and with sample individuals related according to a variety of genealogical configurations, we demonstrated that the top axes of variation from PC-AiR were nearly perfectly correlated with ancestry. In contrast, widely used methods for population structure inference performed poorly in the presence of relatedness, including the PCA method implemented in the EIGENSOFT software, MDS as implemented in PLINK software, and model-based ancestry estimation methods ADMIXTURE and FRAPPE. We also applied PC-AiR and competing methods to the admixed HapMap MXL and ASW population samples. Without using any reference population panels or pedigree information on the sample individuals, the top two axes of variation from PC-AiR nearly perfectly explained proportional European, Native American, and African ancestry in the HapMap MXL and ASW samples as compared to a supervised individual ancestry analysis with ADMIXTURE that included reference population panels. In contrast, all other population structure inference methods were confounded by relatedness, including the FamPCA method which was provided the documented pedigree relationships.

Performing PCA with genome-wide SNP weights that are calculated from external reference panels has recently been proposed (Chen et al., 2013) for certain admixed populations. This approach requires prior knowledge about the ancestries of the individuals in the sample, which may be partially or completely unknown, as well as having available reference panels that are adequate surrogates for ancestry. Nevertheless, the PC-AiR method can also easily incorporate SNP-weights from external reference panels for population structure inference. For example, by designating population samples from external reference panels to be the ancestry representative subset in the PC-AiR algorithm, principal components for individuals in the target sample for population structure inference will be calculated based solely on SNP weights from the reference panels. A potential limitation of using SNP weights from external reference panels, however, is that inference on population structure will be limited to the ancestries of individuals selected from the panels, which may not be representative of the ancestries of all individuals in the sample. An attractive alternative approach would be to perform a PC-AiR analysis on the study sample combined with the external reference panels, where genome-screen data would be used by the algorithm implemented in PC-AiR for the identification of an ancestry representative subset from the combined set of individuals, and where ancestries from both the reference panels and the sample will be allowed to contribute to the SNP weights.

Linear mixed models (LMMs) have recently emerged as a powerful and effective approach for association mapping in samples with population structure as well as family structure or cryptic relatedness (Yang et al., 2014). LMMs have previously been evaluated in samples with subtle population structure (Price et al., 2010; Wu et al., 2011) and have been shown to have appropriate control over type-I error. We evaluated the performance of LMMs in simulation studies where sample individuals have ancestry derived from divergent populations, and our simulation results showed that widely used LMM approaches for association mapping, such as EMMAX and GEMMA, can have an increase in type-I error due to under-correction of SNPs with moderate to high differentiation in allele frequencies between ancestral population, as well as a loss of power due to overcorrection of SNPs with little to no differentiation. This result illustrates potential problems with existing LMM approaches for association mapping in recently admixed populations, where a large proportion of SNPs are expected to have substantial allele frequency differences between the underlying ancestral populations. For example, African Americans have genetic contributions from European and African ancestral populations, and in a comparative analysis of allele frequencies at 1.4 million autosomal SNPs for European (CEU) and West African (YRI) samples in HapMap, we found that approximately 10% of the SNPs were highly differentiated, with allele frequency differences greater than 0.4, while 26% were moderately differentiated, with allele frequency differences between 0.2 and 0.4. Our simulation studies also illustrated that including principal components from PC-AiR as fixed effects in LMMs resulted in appropriate calibration of association test statistics at all SNPs in related admixed samples, protecting against inflated type-I error at highly and moderately differentiated SNPs.

The challenges of inferring genetic ancestry in related samples have been well documented (Patterson et al., 2006; Price et al., 2010). To our knowledge, PC-AiR is the first method to provide robust population structure inference and correction in the presence of known or cryptic relatedness without requiring reference population panels, external SNP loadings, or genealogical information on the sample individuals. We have implemented the PC-AiR method in an R package that is freely downloadable (see Web Resources).

## Acknowledgments

This study was supported in part by the National Institutes of Health grants K01 CA148958 and P01 HG0099568 (to T.T.).

## Disclosure Declaration

The authors have nothing to disclose.

## Web Resources

An implementation of PC-AiR in the R language can be found at: http://faculty.washington.edu/tathornt/software/index.html

## Appendix

Under general population genetic modeling assumptions, we show that for an outbred unrelated pair of individuals *i* and *j*, the 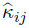 estimator in Equation (1) is a consistent estimator for the quantity

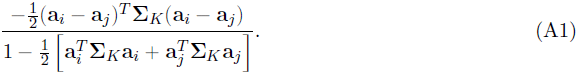

An equivalent expression of the estimator 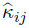 from Equation (1) is

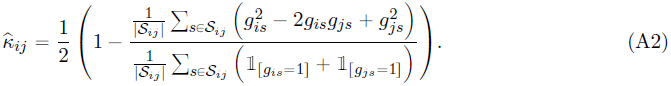

Under our population genetic modeling assumptions, the vector of subpopulation-specific allele frequencies, **p***_s_*, has the properties E[**p***_s_*] = *p_s_***1** and Cov[**p***_s_*] = *p_s_*(1−*p_s_*)**Σ***_K_* for all *s* ∈ *S*. We assume that the ancestral allele frequencies, *p_s_* for *s* ∈ *S*, are independent and identically distributed (i.i.d.) random variables from some unspecified distribution on [0, 1]. Under this assumption, the unconditional expectation of each of the random variables in Equation (A2) is the same for every choice of *s* ∈ *S*, and if we assume that genotypes at different SNPs are independent, then

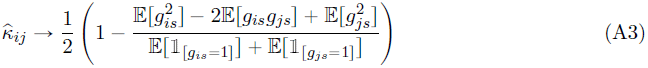

as |*S*| →∞. Note that the independence of SNPs assumption can be relaxed for Equation (A3), and a sufficient condition would be that the effective number of independent SNPs tends to ∞. In what follows, we derive each of the expectations in Equation (A3) conditional on *p_s_*, and we show that the limiting value of 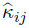 is the value given in Equation (A1), which does not depend on *p_s_*, implying that the i.i.d. assumption can also be relaxed.

Recall that *g_is_* is the number of copies of the reference allele that individual *i* has at SNP *s*, and thus *g_is_* can have a value of 0, 1, or 2. As in Thornton et al. (2012), we define the quantity *µ_is_* to be one half of the expectation of *g_is_*, conditional on individual *i*’s ancestry, **a***_i_*, and the vector of subpopulation-specific allele frequencies, **p***_s_*, at SNP *s*:

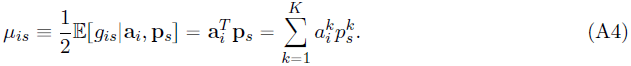

The quantity *µ_is_* can be interpreted as the individual-specific allele frequency for individual *i* at SNP *s*, and it is a linear combination of the subpopulation-specific allele frequencies weighted by individual *i*’s autosomal ancestry proportions from each of the ancestral subpopulations. In Thornton et al. (2012), both **a***_i_* and **p***_s_* are treated as fixed quantities. Here, we similarly treat the ancestry vectors as fixed, and we implicitly condition on **a***_i_* and **a***_j_* throughout what follows, but we allow **p***_s_* to be a random vector for all *s* ∈ *S*. Under our weak basic genetic modeling assumptions, we calculate the following:

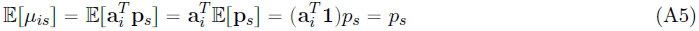

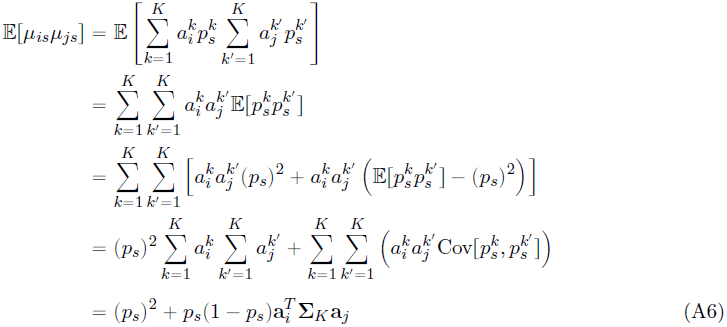

The expectation 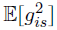 can be obtained directly based on the genotype probabilities for individual *i* conditional on **p***_s_*, where the conditional probabilities that individual *i* has genotype 0, 1, and 2 at SNP *s* are 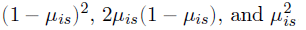, respectively. Thus we obtain

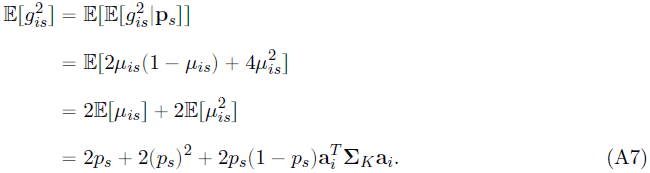

To obtain 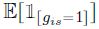, we note that the expectation of an indicator function is just the probability of the event it indicates, so

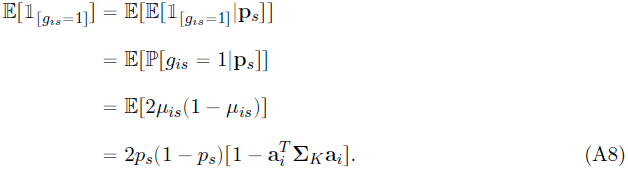

The final expectation that we must obtain for Equation (A3) is E[*g_is_g_js_*]. Since we are only considering unrelated individuals here, the genotype values *g_is_* and *g_js_* are independent conditional on the vector of subpopulation allele frequencies, so 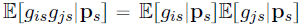. Therefore, we can calculate

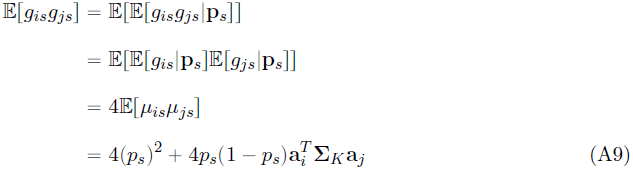

Finally, plugging the expectations given in Equations (A7), (A8), and (A9) into Equation (A3), we have

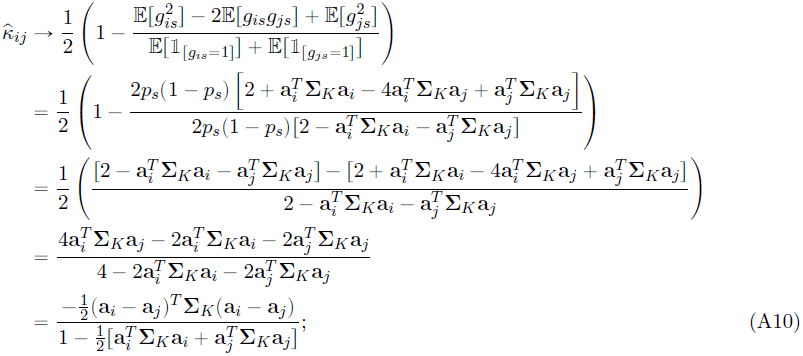

the limiting value of 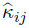 that is given in Equation (A1). Note that the limiting value is not a function of *p_s_*, and thus this convergence holds for any random *p_s_*.

## References

Abney, M., 2009. A graphical algorithm for fast computation of identity coefficients and generalized kinship coefficients. Bioinformatics, 25(12):1561–1563.

Alexander, D. H., Novembre, J., and Lange, K., 2009. Fast model-based estimation of ancestry in unrelated individuals. Genome Research, 19(9):1655–1664.

Balding, D. J. and Nichols, R. A., 1995. A method for quantifying differentiation between populations at multi-allelic loci and its implications for investigating identity and paternity. Genetica, 96:3–12.

Chen, C.-Y., Pollack, S., Hunter, D. J., Hirschhorn, J. N., Kraft, P., and Price, A. L., 2013. Improved ancestry inference using weights from external reference panels. Bioinformatics, 29(11):1399–1406.

Devlin, B. and Roeder, K., 1999. Genomic control for association studies. Biometrics, 55(4):997– 1004.

Gravel, S., Zakharia, F., Moreno-Estrada, A., Byrnes, J. K., Muzzio, M., Rodriguez-Flores, J. L., Kenny, E. E., Gignoux, C. R., Maples, B. K., Guiblet, W., et al., 2013. Reconstructing native american migrations from whole-genome and whole-exome data. PLoS genetics, 9(12):e1004023.

Heath, S. C., Gut, I. G., Brennan, P., McKay, J. D., Bencko, V., Fabianova, E., Foretova, L., Georges, M., Janout, V., Kabesch, M., et al., 2008. Investigation of the fine structure of european populations with applications to disease association studies. European Journal of Human Genetics, 16(12):1413–1429.

International HapMap 3 Consortium, 2010. Integrating common and rare genetic variation in diverse human populations. Nature, 467(7311):52–58.

Kang, H. M., Sul, J. H., Zaitlen, N. A., Kong, S.-y., Freimer, N. B., Sabatti, C., Eskin, E., et al., 2010. Variance component model to account for sample structure in genome-wide association studies. Nature genetics, 42(4):348–354.

Li, J. Z., Absher, D. M., Tang, H., Southwick, A. M., Casto, A. M., Ramachandran, S., Cann, H. M., Barsh, G. S., Feldman, M., Cavalli-Sforza, L. L., et al., 2008. Worldwide human relationships inferred from genome-wide patterns of variation. Science, 319(5866):1100–1104.

Manichaikul, A., Mychaleckyj, J. C., Rich, S. S., Daly, K., Sale, M., and Chen, W.-M., 2010. Robust relationship inference in genome-wide association studies. Bioinformatics, 26(22):2867–2873.

Moltke, I. and Albrechtsen, A., 2014. Relateadmix: a software tool for estimating relatedness between admixed individuals. Bioinformatics, 30(7):1027–1028.

Patterson, N., Price, A. L., and Reich, D., 2006. Population structure and eigenanalysis. PLoS genetics, 2(12):e190.

Price, A. L., Patterson, N. J., Plenge, R. M., Weinblatt, M. E., Shadick, N. A., and Reich, D., 2006. Principal components analysis corrects for stratification in genome-wide association studies. Nature genetics, 38(8):904–909.

Price, A. L., Zaitlen, N. A., Reich, D., and Patterson, N., 2010. New approaches to population stratification in genome-wide association studies. Nature Reviews Genetics, 11(7):459–463.

Pritchard, J. K., Stephens, M., and Donnelly, P., 2000. Inference of population structure using multilocus genotype data. Genetics, 155(2):945–959.

Purcell, S., Neale, B., Todd-Brown, K., Thomas, L., Ferreira, M. A., Bender, D., Maller, J., Sklar, P., De Bakker, P. I., Daly, M. J., et al., 2007. Plink: a tool set for whole-genome association and population-based linkage analyses. The American Journal of Human Genetics, 81(3):559–575.

Staples, J., Nickerson, D. A., and Below, J. E., 2013. Utilizing graph theory to select the largest set of unrelated individuals for genetic analysis. Genetic epidemiology, 37(2):136–141.

Tang, H., Peng, J., Wang, P., and Risch, N. J., 2005. Estimation of individual admixture: analytical and study design considerations. Genetic epidemiology, 28(4):289–301.

Thornton, T., Tang, H., Hoffmann, T. J., Ochs-Balcom, H. M., Caan, B. J., and Risch, N., 2012. Estimating kinship in admixed populations. The American Journal of Human Genetics, 91(1):122– 138.

Thornton, T. A. and Bermejo, J. L., 2014. Local and global ancestry inference and applications to genetic association analysis for admixed populations. Genetic Epidemiology, 38(S1):S5–S12.

Wright, S., 1949. The genetical structure of populations. Annals of eugenics, 15(1):323–354.

Wu, C., DeWan, A., Hoh, J., and Wang, Z., 2011. A comparison of association methods correcting for population stratification in case–control studies. Annals of human genetics, 75(3):418–427.

Yang, J., Benyamin, B., McEvoy, B. P., Gordon, S., Henders, A. K., Nyholt, D. R., Madden, P. A., Heath, A. C., Martin, N. G., Montgomery, G. W., et al., 2010. Common snps explain a large proportion of the heritability for human height. Nature genetics, 42(7):565–569.

Yang, J., Zaitlen, N. A., Goddard, M. E., Visscher, P. M., and Price, A. L., 2014. Advantages and pitfalls in the application of mixed-model association methods. Nature genetics, 46(2):100–106.

Zheng, Q. and Bourgain, C., 2009. Kininbcoef: Calculation of kinship and inbreeding coefficients. (http://galton.uchicago.edu/mcpeek/software/index.html),.

Zhou, X. and Stephens, M., 2012. Genome-wide efficient mixed-model analysis for association studies. Nature genetics, 44(7):821–824.

Zhu, X., Li, S., Cooper, R. S., and Elston, R. C., 2008. A unified association analysis approach for family and unrelated samples correcting for stratification. The American Journal of Human Genetics, 82(2):352–365.

